# Encapsulation of bacteria in bilayer Pluronic thin film hydrogels: a safe format for engineered living materials

**DOI:** 10.1101/2022.09.29.510162

**Authors:** Shardul Bhusari, Juhyun Kim, Karen Polizzi, Shrikrishnan Sankaran, Aránzazu del Campo

**Affiliations:** INM - Leibniz Institute for New Materials, Saarbrücken, Germany; Chemistry Department, Saarland University, 66123 Saarbrücken, Germany; Department of Chemical Engineering, Imperial College London, London SW7 2AZ, United Kingdom; Imperial College Centre for Synthetic Biology, Imperial College London, London SW7 2AZ, United Kingdom; Current affiliation - School of Life Sciences, BK21 FOUR KNU Creative BioResearch Group, Kyungpook National University, Daegu 41566, Republic of Korea

**Keywords:** Engineered living material, bacterial hydrogel, biosensor, bacterial-materials interactions, living therapeutics, biocontainment

## Abstract

In engineered living materials (ELMs) non-living matrices encapsulate microorganisms to acquire capabilities like sensing or biosynthesis. The confinement of the organisms to the matrix and the prevention of overgrowth and escape during the lifetime of the material is necessary for the application of ELMs into real devices. In this study, a bilayer thin film hydrogel of Pluronic F127 and Pluronic F127 acrylate polymers supported on a solid substrate is introduced. The inner hydrogel layer contains genetically engineered bacteria and supports their growth, while the outer layer acts as an envelope and does not allow leakage of the living organisms outside of the film for at least 15 days. Due to the flat and transparent nature of the construct, the thin layer is suited for microscopy and spectroscopy-based analyses. The composition and properties of the inner and outer layer are adjusted independently to fulfil viability and confinement requirements. We demonstrate that bacterial growth and light-induced protein production are possible in the inner layer and their extent is influenced by the crosslinking degree of the used hydrogel. Bacteria inside the hydrogel are viable long term, they can act as lactate-sensors and remain active after storage in phosphate buffer at room temperature for at least 3 weeks. The versatility of bilayer bacteria thin-films is attractive for fundamental studies and for the development of application-oriented ELMs.

## 1. Introduction

Engineered living materials (ELMs) are composites containing microorganisms programmed to perform specific functions. In ELMs for biosensing or drug delivery,^[1][2]^ the organisms-typically bacteria- are encapsulated within hydrogels. These matrices need to support the growth and activity of the organisms, but also biocontainment by preventing the organisms from escaping to the surroundings.^[3][4]^

The growth and the metabolic activity of bacteria embedded in hydrogels depends on the composition and physical properties of the hydrogel matrix.^[3][5]^ In physically crosslinked hydrogels, like agarose or alginate, bacteria grow slower than in free suspension and eventually grow out of the hydrogel (i.e. into the surrounding medium) if there are enough nutrients available.^[6]^ In chemically crosslinked hydrogels, like poly(ethyleneglycol diacrylate) or poly(acrylamide), bacterial growth is restricted to small colonies, independent of the nutrient concentration.^[7]^ Previous studies have demonstrated that Pluronic-based hydrogels are good matrices to encapsulate bacteria.^[3][5][8][9]^ 30 wt. % mixtures of Pluronic F127 and Pluronic F127 diacrylate at different concentrations (named **DA X**, where X represents the fraction of diacrylated Pluronic in the mixture) formed stable hydrogels after covalent crosslinking by photoinitiated radical polymerization of the acrylate groups. The variation of the diacrylate content in the mixture lead to hydrogels with different mechanical properties in terms of elastic moduli (**Table S1**) and viscoelasticity as consequence of the introduction of covalent crosslinks into the physical network.^[5]^ *E. coli* encapsulated in these hydrogels proliferated and kept their metabolic function to an extent which was dependent on the concentration of acrylate groups, i.e. the degree of covalent crosslinks in the hydrogel. In particular, bacteria in DA X hydrogels with X > 50 grew in the form of small, spherical colonies and did not overgrow out of the material^[5]^, suggesting that these could be appropriate materials for a confinement layer. In DA X hydrogels with X<50, bacterial growth was found to be significantly less restricted and controlled.^[5]^ Interestingly, highest rates of inducible protein production were observed in the gels with intermediate degrees of chemical cross-linking (DA X, 25< X <75), indicating that encapsulation influences both growth and metabolic activity of the bacteria in potentially different manner.

While most ELM examples demonstrate bacterial function inside the material, many of them do not address the need for containment.^[9][10]^ This limits the transfer of ELMs into applicable technological solutions. The simplest approach to meet growth and containment requirements for ELMs are bilayer designs, for example core-shell particles or compartments.^[10]^ Tang *et al*. used Ca^2+^ crosslinked alginate as core material and alginate/poly(acrylamide) double network in the shell to encapsulate *E. coli* in beads which retained the organisms for at least 14 days. Liu *et al*. encapsulated liquid cultures of genetically modified *E. coli* in poly(acrylamide)-alginate compartments covered by a polydimethylsiloxane (PDMS) elastomer layer.^[11]^ The resulting living sensor prevented escape of the bacteria for at least 3 days. Connell *et al*. reported compartments made of photo-crosslinked gelatin shells and viscous gelatin cores where bacteria like *Staphylococcus aureus* and *Pseudomonas aeruginosa* could grow and molecularly interact with those in neighboring compartments.^[12]^ Liu *et al*. *f*abricated tattoos using *E. coli* encapsulated in Pluronic hydrogels with a Sylgard 184 (Dow Corning) and Silgel 613 (Wacker) elastomer second layer and demonstrated bacterial retention and function over 8 h.^[13]^ Knierim *et al*. encapsulated *Micrococcus luteus* for gold sequestration and *Nitrobacter winogradskyi* for bioremediation of nitrite in poly(vinylalcohol) fibres and coated it with a poly(p-xylylene) shell by chemical vapor deposition.^[14]^ The shell reduced bacterial escape as a function of the thickness. These examples demonstrate the capabilities and versatility of core-shell designs for bacterial containment in ELMs. In this study, we leverage this design principle to establish a bilayer thin-film platform with which material properties that are optimal for bacterial functions can be investigated while preventing bacterial escape.

The design is based on Pluronic F127 hydrogels. This polymer exhibits a thermo-responsive sol-gel transition at temperatures >14°C at 30 wt. % concentration which results in physically assembled gels. This enables easy fabrication of hydrogel structures involving mixing of the polymer solution with the living organisms at 4°C and the fast gelation at room temperature. In this work, we demonstrate that the combination of a DA X core and a DA100 shell to generate a bilayer thin film is useful for making ELMs wherein bacterial behavior can be tuned by varying the composition of the core while the shell ensures long-term (at least 15 days) biocontainment. The combination of acrylated and non-acrylated polymer allows the design of core and shell out of the same backbone. The use of this format to fabricate live biosensors has been evaluated using engineered L-lactate sensing bacteria relevant for the biotech and biopharma industries.^[15][16][17]^

## 2. Results and Discussion

Bilayer thin films in this work are composed of an inner functional layer containing bacteria embedded in 30 wt. % DA X hydrogel, which is sandwiched between a containment layer of 30 wt. % DA 100 and a glass slide **(Figure 1)**. The bilayers were fabricated manually by pipetting the solutions at 0 °C onto the glass slide (functional layer) and onto a releasable parafilm (containment layer), allowing the physical gel to form at room temperature, and photopolymerizing the acrylate end-groups in both layers in a single exposure step. (Figure 1a) The initial volumes of the inner and outer layers were 2 and 30 μL. The outer diameter of the thin film was ca. 11 mm and the inner diameter was 2-3 mm. Bilayers with different DA X compositions of the inner layer were prepared in order to identify the best possible combination that maximizes the metabolic activity of the bacteria. The thin films were transparent and allowed microscopy imaging of the bacteria in the inner gel.

**Figure 1.**
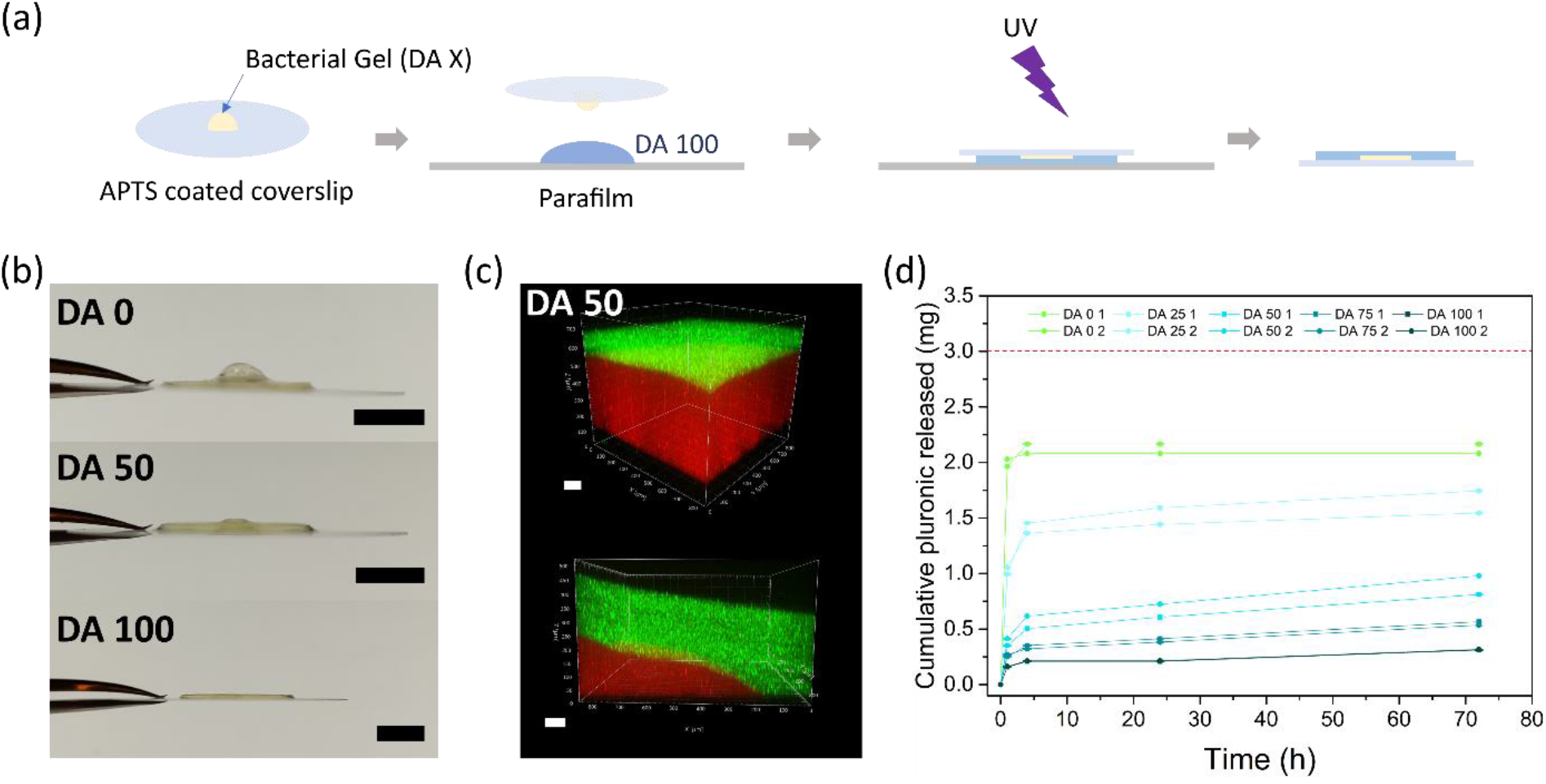
Bilayer thin films prepared with DA X inner layers and a DA 100 outer layer. **a)** Fabrication steps to obtain the bilayer thin films. A drop of DA X/bacteria mixture is placed on an acryl-terminated coverslip and pressed onto a larger DA 100 drop on a parafilm. The drops are pipetted from solutions on ice and the bilayer is left at room temperature to form the physical gel. UV exposure activates the covalent crosslinking of the two layers; **b)** Side view images of a DA 0, DA 50 and DA 100 bilayer thin films with bacterial inner gels at 24 h showing the central protrusion as consequence of higher swelling in the inner gel (Scale = 5 mm); **c)** Confocal microscopy imaging of a swollen bilayer thin film with DA 50 inner core. Z-stack images were taken in the middle of the inner gel and at the edge of the inner-outer layer interface after swelling for 24 h. DA 50 inner gel contains red fluorescent beads and DA 100 outer gel contains green fluorescent beads (Scale = 100 μm) and **d)** Cumulative release profile of Pluronic molecules from DA X hydrogel discs during 3 days in water as calculated from gel permeation chromatography (GPC) measurements (N = 2 and n ≥ 3, whiskers indicate mean ± standard deviation). The maximum theoretical Pluronic concentration that can be released from the films according to the mass of the thin film is 3 mg (dotted red line).

When immersed in water or buffer solutions, DA 0 hydrogels swell and dissolve since they are only held by physical interactions. Müller *et al*. showed a 382% swelling degree for DA 15 discs.^[18]^ The swelling degree of DA X films in our study decreases with increasing X value (i.e. increasing covalent crosslinking, **Figure S1**) in films with X ≥ 25. The DA X films retain their format after swelling. In the case of the bilayers, swelling results in a protrusion in the middle of the bilayer as consequence of the higher swelling of the DA 0-75 inner layers with lower crosslinking degree than the outer DA 100 shell (Figure 1b-c, **S2**). The thickness of the outer layer after swelling is ca. 100-150 μm (measured on the top of the protrusion, Figure 1c).

Swelling of the Pluronic hydrogels is accompanied by leakage of non-diacrylated Pluronic from the bilayer into the water phase. This could be an advantageous feature for bilayer-based ELMs, since it increases the porosity of the hydrogel after film formation and swelling, and this can favor diffusion of nutrients and cell viability.^[18]^ We tested the release of Pluronic molecules from DA X hydrogel films after immersing them in water for up to 72 hours by GPC analysis of the supernatant. Pluronic was released to the medium from all DA X hydrogels within 1-2 hours (Figure 1d, **S3**). At longer time scales, slow release was observed in DA 25-75 hydrogels and no release in DA 100. The molar mass distribution of the released polymer corresponded to that of the Pluronic precursor, indicating that only molecules that are not crosslinked were able to diffuse out of the hydrogel. The amount of pluronic released into the supernatant decreased with increasing PluDA content, from ca. 55 % of the film mass in DA 25 to ca. 10 % of the film mass in DA 100 after 72 h. Note that the degree of substitution of PluDA was 70 %, meaning that all hydrogels contain at least 30 % of Pluronic chains with only one or no acrylated chains. Although DA 0 samples dissolved completely, the amount of Pluronic detected in the supernatant was lower than the expected polymer mass in the film (represented by the red dotted line). This could be due to (i) adsorption of Pluronic molecules on the surface of the wells and pipettes or (ii) micellar association of Pluronic molecules in the solution.

To assess the behavior of encapsulated bacteria, we fabricated bilayer thin films containing *E. coli* in inner layer hydrogels made with different DA X compositions. Brightfield and fluorescence microscopy images of live/dead stained bacteria after 24 h revealed that bacteria grew in all the inner gels, although with different morphologies in the different composition **(Figure 2)**. In DA 0, i.e. inner layer with no chemical crosslinking, the bacteria grew as a uniform biomass, suggesting that the gel network did not constrain the growth of the organisms. Despite swelling of the inner layer causing the outer DA 100 layer to expand, bacterial containment was not compromised within the timescale of the experiment. In all other compositions, bacteria formed colonies with individual, elongated (DA 25) or rounded (DA 50 – DA 100) geometries visible under the microscope. Such morphologies are in line with the results of bacteria growth within DA X hydrogels within closed microchannels from our previous report.^[5]^ In DA25 hydrogels colonies showed an elongated morphology in the vertical direction, towards the supernatant. This could be a consequence of mechanical effects during swelling. Note that in thin films the hydrogels are exposed to medium and there is a release of Pluronic during the first hours. This indicates that the covalently cross-linked DA X network restricts growth also after swelling and loss of polymer fraction. No bacterial motility was observed in the inner gel for all constructs except for DA 0. The staining further revealed that most of the bacteria were alive, suggesting that the bilayer sustains bacteria function.

**Figure 2.**
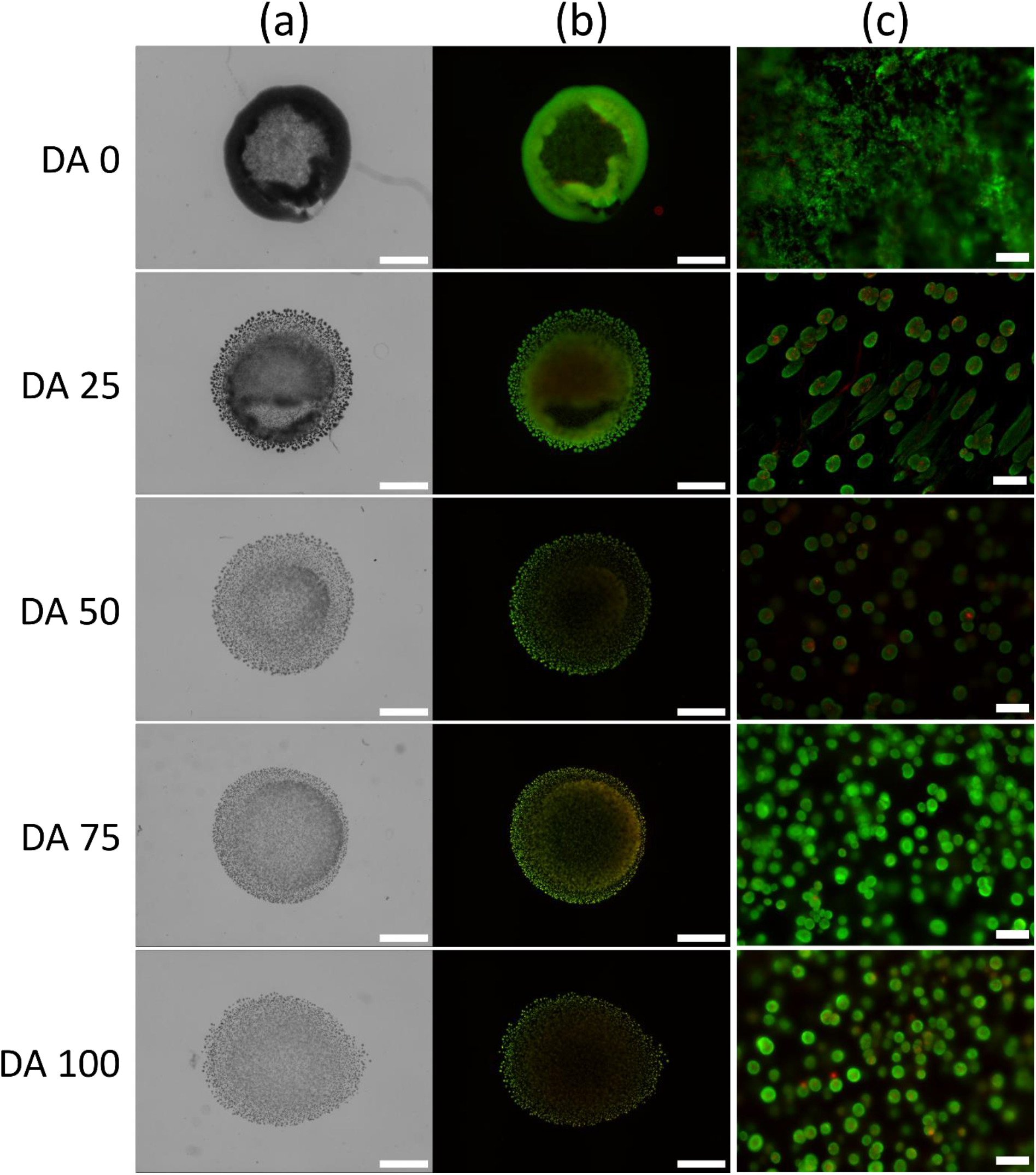
Viability of bacterial populations within the bilayer thin films. **a)** Brightfield and **b)** fluorescence microscopy images showing *E. coli* growth within bilayer thin films with different DA X inner layer. Images were taken near the bottom of the inner gel. (scale bar = 1000 μm); **c)**Representative images of bacterial colonies at a higher magnification after 24 h. (scale = 50 μm) Samples were stained with SYTO 9 (green = live) and Propidium Iodide (red = dead) after 24 h of incubation.

In ELMs developed as biosensors or therapeutic devices, function is often associated to production of a protein in response to an external stimulus or inducer molecule. Inducible protein production in our thin films was tested using an optogenetically engineered *E. coli* strain that expresses a red fluorescent protein (RFP) in response to blue light.^[5][6]^ With pulsed light irradiation (470 nm, 500 ms ON, 10 min OFF), fluorescence was detectable in the inner film within 4 hours, although it was not homogeneous through the film. Higher fluorescence intensity was observed at the outer edge of the inner layer (**Figure 3a**). This could be a consequence of a higher nutrient concentration, which would lead to a higher bacterial density^[19]^. We estimated the overall kinetics of protein production across the film by quantifying the total fluorescence intensity for 18 hours. An increase in fluorescence was observed during the first 4-10 hours (**Figure 3b**) and a plateau value was reached at longer times. The plateau was reached when fluorescence intensity exceeded the fluorescence saturation level of the microscope required to observe the early stage of RFP production. Therefore, it does not reflect the real RFP concentration in the film at longer time scales. We analyzed the slope of the linear part or the fluorescence curves for an estimation of the kinetics of RFP expression in the different systems. The result of this analysis is shown in **Figure 3c** and reveals faster RFP production in DA 50 (5.1 ± 0.1 au/h) compared to the other compositions (DA 0 = 3.1 ± 0.1 au/h, DA 25 = 4.8 ± 0.1 au/h, DA75 = 4.8 ± 0.1 au/h, DA 100 = 4.0 ± 0.1 au/h). RFP production in DA 50 inner layers also showed the smallest deviation across the whole period of the experiment. The higher rate of protein expression in DA 50 hydrogels is in agreement with our previous observations of RFP production in bacterial DA X hydrogels within closed microchannels.^[5]^

**Figure 3.**
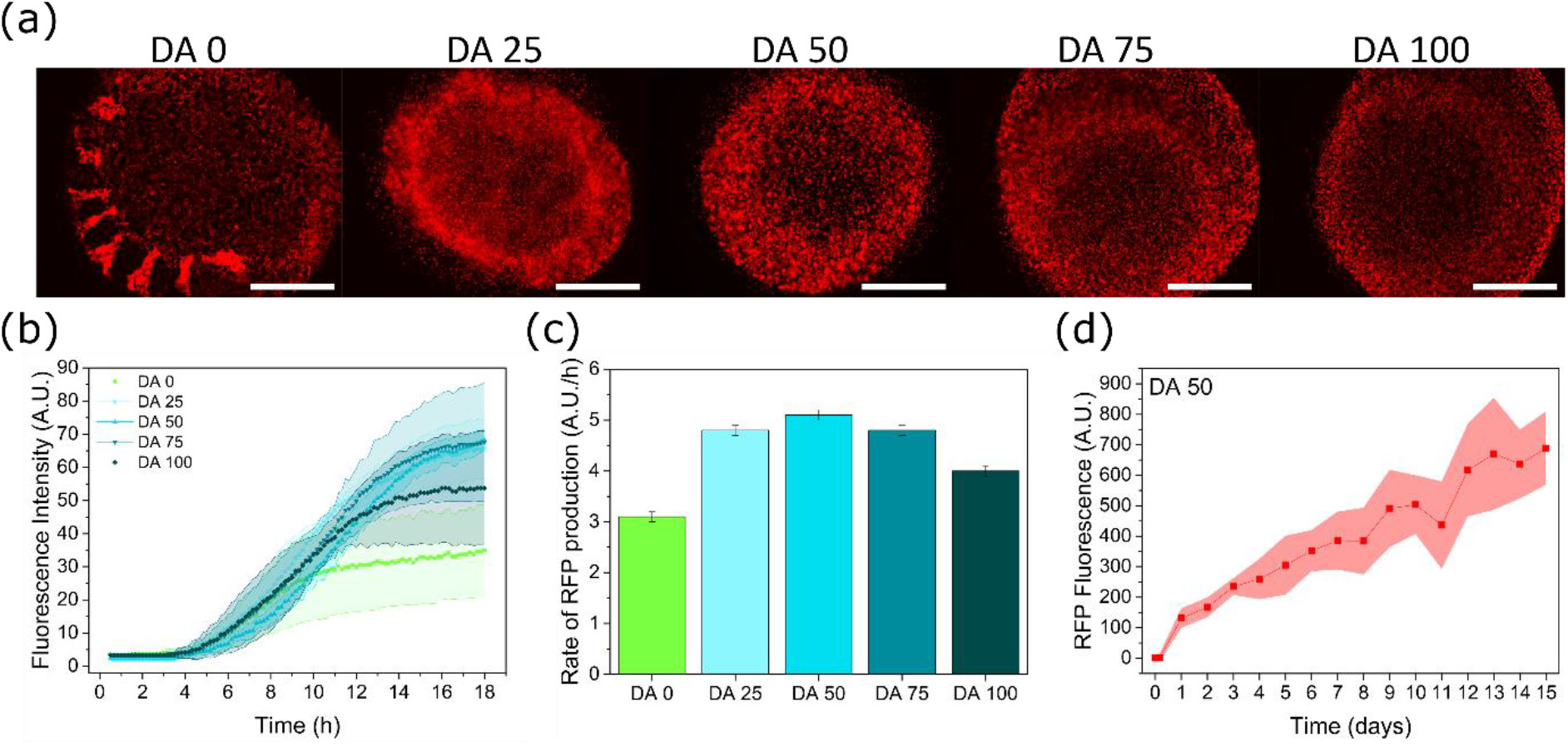
RFP production within the bilayer thin films. **a)** Fluorescence images of RFP producing bacterial gels indicating RFP expressed by the encapsulated bacterial colonies at 12 h (scale = 1000 μm); **b)** Quantification of total fluorescence intensity of the bilayer thin films during 24 h from fluorescence microscope images in (a) (N=3, mean ± standard deviation); **c)** Rate of RFP production in bilayers with different DA X layer compositions estimated from the slope of the linear part of the fluorescence intensity curves in (b) (mean ± standard deviation) and **d)** Quantification of fluorescence intensity within DA 50 bacterial gel thin films for 15 days using the plate reader (N=8 until day 12, N=4 until day 15, mean ± standard deviation, medium was changed every 2 days).

We selected DA 50 as inner hydrogel for the next experiments and tested the safe encapsulation of the bacteria in the film and the activity of the bacterial bilayers over longer periods of time (Figure 3d). Bilayer thin films were cultured for over two weeks. No outgrowth of bacteria was detected in the biofilms incubated for over 10 days (**Figure S4**), demonstrating the capability of bilayer hydrogels with DA 50 inner layer and DA 100 outer layer to safely support activity while confining bacteria colonies. Beyond 15 days, loss of adhesion between the gel and the glass substrate was observed in half of the samples, indicating that the stability of the thin film is not limited by the stability of the enveloping/outer hydrogel, but rather by the design of the glass-hydrogel interface. Optimization of the glass treatment step could be a simple solution for longer studies. The overall fluorescence signal of the bilayer increased during 15 days. Note that this quantification was done using a microplate reader that provided a larger dynamic range for measuring fluorescence intensity than the microscope images in Figure 3a. Analysis of the supernatant indicated that RFP was released from the bilayer hydrogels already on day 2, and at a higher concentration on day 8 (**Figure S5**).^[6]^ In this experiment the medium was changed every two days. Since the bacterial colonies are not expected to considerably increase in number or size beyond 1 day (**Figure S6**), the increase in fluorescence suggests that the protein is continuously secreted and accumulates within the network.

We then tested the suitability of the bilayer hydrogels for biosensing by encapsulating bacteria engineered as L-lactate biosensors. Lactate biosensors are used to monitor the health of mammalian cell cultures, for which secure encapsulation would be a fundamental requirement.^[17]^ This metabolite is a waste product in mammalian cell cultures formed as a byproduct of glycolysis. High concentration of lactate causes acidification of the medium and impacts growth and productivity in the culture.^[20]^ Current lactate monitoring is done by HPLC or electrochemical analysis methods, which have limits in terms of cost, sensitivity, and robustness. Industry standard instruments (eg. Bioprofile Analyzer) and kits for lactate sensing based on enzymatic reactions have high costs associated with the purified enzymes. To address these issues, bacterial biosensors able to sense lactate in mammalian cell culture medium and report it by means of a fluorescent protein have been developed.^[15][16][17]^ To be used in combination with biotech reactors, bacteria were encapsulated within giant unilammellar vesicles^[16]^ or multilayer polymer shells^[17]^. These methods ensured confinement of the bacteria on short timescales (24 h or less), but the long-term behavior was not examined. Thus, we encapsulated the living L-lactate biosensors in DA 50 /DA 100 bilayers and tested their functionality.

To ensure that variability in medium oxygen concentrations do not affect the sensing performance, the bacteria were engineered to produce an oxygen-independent fluorescent protein, CreiLOV, as the reporter. Although this fluorescent protein shows lower fluorescence intensity and quicker bleaching than the GFP used in previous studies,^[15][16][17]^ the signal was high enough to be observed within the gels. In line with previous reports of encapsulated lactate sensing bacteria^[17]^, fluorescence signals in response to 1 mM and 10 mM L-lactate were observed after 9 and 6 h, respectively (**Figure 4a**). This slow response might be due to the dependence of fluorescent protein production on bacterial growth. In a second experiment, bacteria were allowed to grow inside the gels for 6h after which L-lactate was added (Figure 4b). An increase in the fluorescence signal with 10 mM lactate was observed already within the first 2 h, supporting the view that protein production is growth-dependent, which is in line with our previous study.^[5]^ However, a higher degree of leaky expression was also observed in the absence of L-lactate, which overlapped with the signal generated by 1 mM lactate. This leaky expression of CreiLOV could be caused by incomplete repression of the promoter by the lldR repressor. The repressor expression is driven by a medium-strength constitutive promoter (BBa_J23118)^[21]^ optimized for lactate-switchable repression in rapidly growing bacterial cultures. Once expressed, the main mechanism for removal of CreiLOV is by cell division. Therefore, small amounts of leaky expression can accumulate a high signal when the bacterial growth is restricted under encapsulation conditions. This effect is also observed when the bacterial biosensors are grown on agar plates as opposed to growth in culture (**Figure S7**). This highlights the importance of understanding the unique characteristics of bacterial growth and functionality in confinement. Studies are underway to reduce the leaky expression of CreiLOV either through stronger expression of the repressor or through degradation of the reporter in the absence of L-lactate.

**Figure 4.**
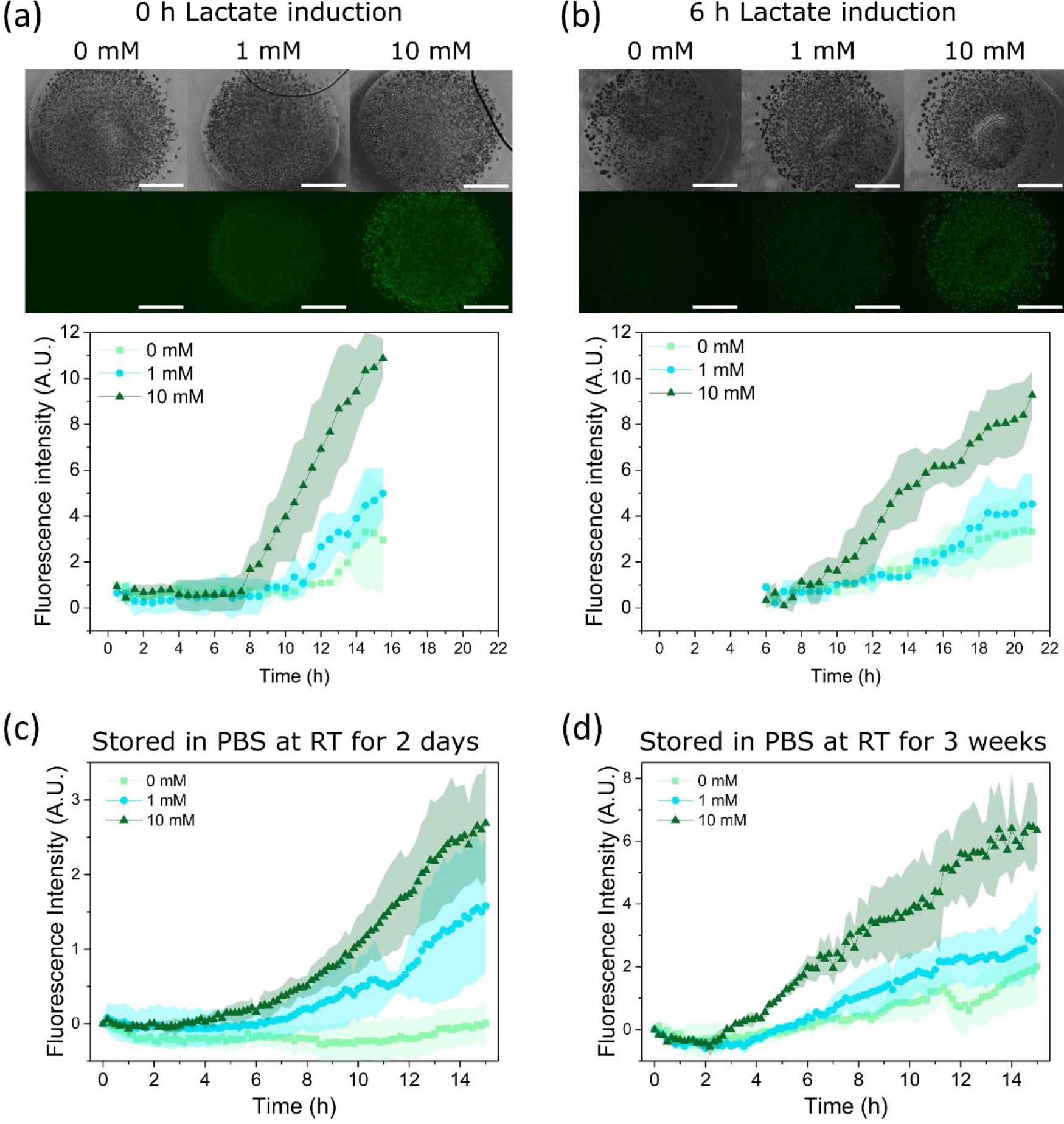
Lactate inducible creiLOV production within the bilayer thin films. Brightfield and fluorescence microscopy images at 15:30 h (scale = 1000 μm) and quantification of fluorescence intensity within DA 50 bacterial gel thin films upon lactate induction at **a)** 0 h (N=2) and **b)** 6 h (N=3) indicating creiLOV production (mean ± standard deviation); Quantification of fluorescence intensity within DA 50 bacterial gel thin films upon lactate induction post growth for 1 day and storage in PBS for **c)** 2 days (N=2 for 0 and 1 mM, N=3 for 10 mM) and **d)** 3 weeks (N=3) indicating creiLOV production during 15:30 h (mean ± standard deviation).

The possibility to store the bilayer thin-film sensor units at room temperature was tested. The bacteria in the hydrogels were allowed to grow for a day and the bilayer thin films were stored in phosphate buffered saline (PBS) for 2 days or 3 weeks (Figure 4c and 4d), after which they were transferred to cell-culture medium containing L-lactate. The sensing capability and response rate of the sensor units remained unchanged: fluorescence increase was observed at between 2 - 4 h after addition of 10 mM L-lactate. Leaky expression in the absence of L-lactate was also observed to the same degree as with the 6 h incubation condition. Once again, this level of leaky expression overlapped with the response to 1 mM L-lactate, although the two conditions could be distinguished beyond 8 h. As a storage alternative, freeze drying was tested (**Figure S8**). On reconstitution of the sensor units with medium containing L-lactate, fluorescence signals were observed to increase at around 4 h. However, freeze-drying damaged some of the samples and bacteria escaped into the surrounding medium. The fluorescence readouts from these samples were measured spectrophotometrically using a multiplate reader, indicating the possibility to develop these sensors for real-time sensing in a bioreactor using fluorimeters, similar to how sensing pads are currently used for monitoring O2, CO_2_ and pH in culture flasks.^[22][23][24]^

## 3. Conclusions

This study describes a secure bacterial encapsulation format using Pluronic F127 based hydrogels for creating ELMs. Using hydrogel bilayers, the properties of the inner bacterial layer can be adjusted to maximize growth and metabolic function, while the outer, highly chemically cross-linked layer, is designed to prevent leakage of cells. These layers are polymerized onto a glass cover slip in one step and result in a transparent thin film that enables the use of this format for microscopy and spectroscopy analysis of bacterial behavior inside. We show that these hydrogels support bacterial growth and protein production. The highest rate of inducible protein production was observed in bilayers with DA 50 inner layer and could be maintained for at least 2 weeks. The produced RFP was released from within the gel from day 2 and increased in following days, indicating that the hydrogels are permeable and sustain diffusion of the produced molecules long term. By encapsulating L-lactate sensing bacteria, we illustrated the possible application of bacterial bilayers as biosensors. Although the leaky expression of the protein needs to be addressed in the lactate sensing genetic circuit, our results show how the performance and safety of ELMs, which is crucial for application purposes, can be improved with dedicated but simple designs.

## 4. Materials and Methods

### 4.1. Bacteria cultures

The endotoxin free strain of *E. coli* (ClearColi(R) BL21(DE3), BioCat)^[6]^ was used for the bacterial studies. It was transformed with the plasmid pUC19 to enable Ampicillin resistance and minimize the risk of contamination in the culture. Transformation in ClearColi was performed by electroporation as described by the manufacturer of these competent cells. Bacterial cultures were grown for 16 h at 35°C, 180 rpm in LB Miller medium supplemented with NaCl and 50 μg/mL of Ampicillin to an optical density at 600 nm wavelength (OD600) between 0.5 – 1. For the fluorescence based experiments, we used a previously optogenetically engineered strain of ClearColi that produces red fluorescence protein (RFP) when illuminated with blue light.^[6]^ The RFP producing strain harbors the plasmid pDawn-RFP that encodes blue-light activatable gene expression and provides kanamycin resistance. The red fluorescence signal was used to quantify protein production and the colony volume measurements in hydrogel-encapsulated bacterial populations. The bacteria were cultured for 16h at 35°C, 220 rpm in LB Miller medium supplemented with NaCl and 50 μg/mL of Kanamycin to an optical density at 600 nm wavelength (OD600) value between 0.5 – 1.5. All procedures with the light-inducible bacterial strain were performed either in the dark or under a laminar hood with an orange film that cuts off blue light to prevent their pre-mature induction.

### 4.2. Construction of the lactate biosensor

A plasmid of the lactate biosensor plasmid^[17]^ derivative containing CreiLOV reporter was used to develop lactate biosensor for low oxygen conditions. *E. coli* DH5α cells were used for expression and characterization of all the plasmids constructs. Briefly, the reporter module carrying the lldR promoter and CreiLOV gene was digested with EcoRI and SacI and cloned into the same sites of the biosensor plasmid.^[17]^ The new construction was propagated in BW25113 (ΔArcA) strain, from the KEIO collection, to avoid repression of aerobic pathways.^[25]^ Bacterial cultures were grown for 16 h at 35°C, 180 rpm in LB Miller medium supplemented with 50 μg/mL of kanamycin to an optical density at 600 nm wavelength (OD600) between 0.5 – 1.

### 4.3. Functionalization of coverslips with (3-acryloxypropyl)trimethoxysilane to improve attachment of the bilayer hydrogel to the supporting glass

13/25 mm glass coverslips were sonicated in ethanol for 5 min and then rinsed with ethanol. The coverslips were treated with 1% v/v solution of (3-acryloxypropyl)trimethoxysilane (APTS, Merck - Sigma Aldrich) in ethanol for 1 h. The coverslips were then rinsed in ethanol and dried for further use.

### 4.4. Preparation of bacteria-DA X thin films

Pluronic F127 (Plu, MW~12600 g/mol, Sigma-Aldrich) and Pluronic F127 diacrylate (PluDA) polymers with substitution degree of 70% were used for the studies. The synthesis of PluDA has been reported elswhere.^[26]^ DA 0 and DA 100 precursor solutions were prepared by diluting Plu or PluDA at 30% (w/v) in milliQ water contining 0.2% w/v Irgacure 2959 photoinitiator. DA 25, DA 50 and DA 75 precursor solutions were prepared from mixtures of DA 0 or DA 100 solutions in ratios as shown in **Table 1**. For the inner hydrogel with bacteria, precursor solutions at 4°C were added to the bacterial suspension (OD600 of 0.5, ~ 4 × 10^7^ cells/mL) at 9/1 (v/v) ratio to achieve a final OD600 of 0.05. The bacterial mixture was vortexed for 30 s to homogeneously disperse the bacteria.

To fabricate the thin films, 2 μL of the inner hydrogel precursor mixture containing the bacteria was placed on a coverslip previously functionalized with APTS and left at room temperature for 10 min for gelation. This step introduces acryl groups at the surface of the glass slide which react with DA X precursor solutions during the photopolymerization step.^[27][28]^ This step covalently anchors DA X hydrogels to the glass slide and prevents detachment of the thin film from the glass slide during the experiment. To fabricate the DA100 outer layer, 30 μL of DA 100 precursor solution was dropped on parafilm and allowed to physically crosslink at room temperature for 10 min. The coverslip with the bacterial gel was pressed against the DA 100 gel on the parafilm. The hydrogel sandwich was kept at room temperature for 10 min for physically crosslinked gelation to occur and exposed for 60 s to a UV Lamp inside Alpha Innotech FluorChem Q system (Biozym, Oldendorf, Germany) (6 mW / cm^2^) which was the illumination step used to initiate the photopolymerization of the acrylate groups and covalent crosslinking of the hydrogels. After additional 10 min, the parafilm was peeled off. The thin-film construct was incubated in medium at 37°C with 5% CO_2_.

### 4.5. Quantification of swelling ratio

DA X hydrogels discs (20 μL polymer precursor solution) were prepared on APTS coated glass coverslips using PDMS molds of diameter of 6 mm and height of 0.8 mm. The samples were irradiated with UV for 60 s, same as mentioned above. The hydrogels were immersed in 500 μL of MQ water and incubated at 37 °C and 5 % CO_2_. The swelling ratios were gravimetrically determined in duplicate by dividing the wet hydrogel weight at timepoint t, Wt, by the initial hydrogel weight, Wi:

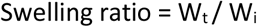

Samples were weighed at specific time points (1, 2, 3 and 24 h) and the supernatant was replaced with fresh MQ water at each timepoint.

### 4.6. GPC analysis of Pluronic in supernatant

DA X hydrogels films (10 μL polymer precursor solution, ca. 3 mg of polymer) were prepared on APTS coated glass coverslips by inverting them on different DA X hydrogels on parafilm and irradiated with UV. The hydrogels were immersed in 500 μL of MQ water and incubated at 37 °C and 5 % CO_2_. At specific time points (10 s, 1 h, 4 h, 24 h and 72 h), the supernatant was taken out from the plate and immediately replaced with fresh MQ water. For the GPC measurement, PSS GPC-MALLS System was used. Injection volume was 100 μl with the mobile phase comprising of 100 % 0.5 g/L NaN3 and 15 % ACN. Flow rate was 1 ml/min. A PSS SUPREMA LUX separation column was used (dimensions 8 x 50mm^2^, particle size 10 μm) and a PSS SUPREMA Linear M (dimensions 8 x 300 mm^2^, particle size 10 μm). The measurements were performed at 25 °C and 37 bars pressure. An Agilent detector with positive signal polarity was used at temperature of 25 °C. Each sample at 10s, 4 h, 24 h and 72 h was measured four times and samples at 1 h were measured three times. The concentration of the released pluronic in the supernatant was determined using a standard calibration curve of known pluronic concentrations (**Figure S3**).

### 4.7. Bilayer thin film thickness

For visualising the bilayers, 0.2 μm FluoSpheres Carboxylate-Modified Microspheres (Invitrogen) were used at 0.02 % concentration within the DA 100 outer gel (yellow green) and DA 50 inner gel (crimson colored). The thin film was then imaged using Zeiss LSM 880 confocal laser scanning microscopy at 24 h using EC Plan-Neofluar 10×/0.30 M27 objective. The green and red beads were excited using 488 nm and 633 nm laser and the emission spectra was detected at wavelength of 493-628 nm and 638-755 nm respectively. Z-stacks were taken in a z-step size of ca. 5.97 μm. Imaris software (Version 9.8.2, Bitplane, Zurich, Switzerland) was used to process the z-stacks to create 3D images.

### 4.8. Staining and imaging of bacterial colonies within thin films

The LIVE/DEAD BacLight Bacterial Viability kit (Thermo Fisher Scientific L7012) was used to visualize growth of bacterial colonies. Stock solutions of the stain were stored at −20°C. To make the working solution, 3 μL of the stock solutions were added to 1 mL of phosphate buffered saline (PBS). The bacterial hydrogels were washed with PBS to remove traces of the medium and incubated with 100 μL of the stain solution for 30 min. The samples were washed with 0.5 mL PBS once and imaged under the microscope. For fluorescence imaging, BZ-X Filter TRITC OP-87764 (excitation 545/25, emission 605/70) was used for imaging the dead colonies while BZ-X Filter GFP OP-87763 (excitation 470/40, emission 525/50) was used for imaging the live colonies using the Keyence PlanFluor 4×/0.13 objective. Nikon Ti-Ecllipse (Nikon Instruments Europe B.V., Germany) microscope with S Plan Fluor ELWD 20x/0.45 objective used the Semrock filters (Semrock Inc., Rochester, USA) (LF488-C-000 filter, excitation 482/18, emission 525/45 for live colonies and LF561-B-000 filter, excitation 561/14, emission 609/54 for dead colonies). Image processing and analyses were performed using Fiji edition of ImageJ (ImageJ Java 1.8.0).

### 4.9. Analysis of RFP production by bacteria in bilayer thin film format

Bilayer thin films containing bacteria in the inner gels were freshly prepared and 400 μL of LB Miller medium supplemented with NaCl and 50 μg/mL of Kanamycin was added. RFP production was activated at 0-0.5 h by illuminating the thin films with blue light (BZ-X Filter GFP OP-87763, excitation 470/40, emission 525/50) pulses of 500 ms every 10 min for 24 h using the Keyence PlanFluor 4×/0.13 objective. The thin films were kept in static conditions at 37 °C and 5 % CO_2_ using an incubation chamber (Stage Top Incubator, Tokai Hit) coupled to a BZ-X800 (Keyence, Osaka, Japan) microscope. For fluorescence imaging to detect the RFP production, BZ-X Filter TRITC OP-87764 (excitation 545/25, emission 605/70) was used. The light intensity and exposure settings were optimized for detecting early time-point generation of fluorescence and following increase in intensity for several hours without exceeding saturation limits. Image processing and analyses were performed using Fiji edition of ImageJ (ImageJ Java 1.8.0). Quantification of the overall fluorescence intensity of the film was done by determining the mean grey value post thresholding (Mean autothreshold) the colonies.

### 4.10. Long term stability and RFP production from bilayer films

For the long-term (2 weeks) studies of the stability and RFP production of the bacteria hydrogels, the bilayer thin film with DA 50 inner layer were stored in 400 μL of LB Miller medium supplemented with NaCl and 50 μg/mL of Kanamycin. The medium was changed every two days. The biofilms were illuminated with blue light pulses of 2 s every minute. The RFP protein production was measured using a black glass bottom 24 well plate in a plate reader (Tecan Infinite M200-Pro, excitation 545 nm, emission 605 nm) using the setting of multiple reads per well. A matrix of 3 × 3 reads were taken within the 24 well plate and the position in the middle of the plate, where the inner gel of the thin film was placed, was considered for the measurements. The growth of fluorescence was quantified with respect to the reading at 0 h. The absence of leakage of the thin films was observed under the microscope and confirmed by streaking 4 μL of the supernatants (day 10) on agar plates (made up of LB supplemented with NaCl and 50 μg/mL of Kanamycin) and incubated at 28 °C for 2 days. No colonies were seen on the agar plate. (**Figure S4**) To quantify RFP production, Strep-Tactin XT affinity beads (IBA Life Sciences) were spun down and washed twice with PBS at 300 rpm for 60 s. 20 ul of the beads were added to 50 ul of the supernatant solution and incubated for 1 h. The mixture solution was then pipetted in a 24-well plate and the beads were imaged using Nikon Ti-Ecllipse (Nikon Instruments Europe B.V., Germany) microscope with SPlanFluor 20x LWD/ 0.7 objective using the Semrock LF561-B-000 filter (excitation 561/14, emission 609/54).

For the brightfield microscopy analysis over 3 days, Nikon Ti-Ecllipse (Nikon Instruments Europe B.V., Germany) microscope with Plan Apo λ 10×/0.45. Thin films bacterial hydrogels were incubated at 37 °C and 5 % CO_2_ using an Okolab incubation chamber (Okolab SRL, Pozzuoli, Italy) coupled to the microscope. Imaging locations were selected near the middle of the bacterial gel.

### 4.11. L-lactate biosensors

For the biosensing platform, BZ-X Filter GFP OP-87763 (excitation 470/40, emission 525/50) and Keyence PlanFluor 4×/0.13 objective were used to detect the production of creiLOV protein. The thin films were incubated in an incubation chamber (Stage Top Incubator, Tokai Hit) coupled to a BZ-X800 (Keyence, Osaka, Japan) microscope in 400 μL of RPMI 1640 (Thermo Fisher Scientific, Germany) medium supplemented with 50 μg/mL of Kanamycin to lower the background signal which was seen in case of LB medium. L-lactate solution in phosphate saline buffer was added in desired concentrations (0, 1 and 10 mM) to RPMI 1640 medium added to the samples. L-lactate was added to the samples at 0 h/ 6 h timepoints followed by incubation at 37 °C and 5 % CO_2_ during which creiLOV fluorescence images were taken every 30 min and quantified by determining the mean grey value post thresholding (Mean autothreshold) the colonies and subtracting the mean grey value of the background. To test the storage efficacy of this living biosensors platform, the thin films were incubated at 37 °C and 5 % CO_2_ to allow the bacteria to grow for 1 day and then stored in PBS for 2 days at room temperature. L-lactate was added and the development of the creiLOV fluorescence was then recorded in a TECAN plate reader every 15 min (excitation 447 nm, emission 497 nm) using the setting of multiple reads per well, as described earlier. For the freeze-drying experiments, the thin films were first soaked in 12 % w/v of sucrose solution for 1 day post growth of 1 day, dipped directly into liquid nitrogen for 2 min and freeze-dried using the freeze dryer (Christ Alpha 1-2 LDplus, Germany) at −20 °C at 1 mbar for 5 h. L-lactate was added to freshly added RPMI medium and the development of the creiLOV fluorescence was recorded as above.

### 4.12. Statistical analyses

One-way analysis of variance (ANOVA) with post hoc Tukey HSD test was performed with results involving more than three data points. Differences were considered statistically significant at * p < 0.05, ** p < 0.01, *** p < 0.001. Analyses were performed with Origin Pro 9.1 software.

## Acknowledgements

SB, SS and AdC acknowledge funding support from the Collaborative Research Centre SFB 1027 (Deutsche Forschungsgemeinschaft) and from the Leibniz ScienceCampus Living Therapeutic Materials, LifeMat (W32/2019, Leibniz Gemeinschaft). Authors thank Dr. Mitchell Han and Dr. Cao Nguyen Duong for insights in image analysis, Ana Díaz Álvarez for functionalization of glass coverslips, Dr. Claudia Fink-Straube and Thi Vinh Ha Rimbach-Nguyen for gel permeation chromatography (GPC) measurements and Dr. Samuel Pearson for his support in the interpretation of GPC results.

## References

[1] L. Xu, X. Wang, F. Sun, Y. Cao, C. Zhong, W.-B. Zhang, Current Opinion in Solid State and Materials Science 2021, 25.

[2] A. Rodrigo-Navarro, S. Sankaran, M. J. Dalby, A. del Campo, M. Salmeron-Sanchez, Nature Reviews Materials 2021, 6, 1175.

[3] H. Priks, T. Butelmann, A. Illarionov, T. G. Johnston, C. Fellin, T. Tamm, A. Nelson, R. Kumar, P. J. Lahtvee, ACS Applied Bio Materials 2020, 3, 4273.

[4] M. Schaffner, P. A. Rühs, F. Coulter, S. Kilcher, A. R. Studart, Science Advances 2017, 3, DOI 10.1126/sciadv.aao6804.

[5] S. Bhusari, S. Sankaran, A. del Campo, Advanced Science 2022, 2106026, 1.

[6] S. Sankaran, A. Campo, Advanced Biosystems 2019, 3, 1800312.

[7] S. Sankaran, J. Becker, C. Wittmann, A. Campo, Small 2019, 15, 1804717.

[8] A. Saha, T. G. Johnston, R. T. Shafranek, C. J. Goodman, J. G. Zalatan, D. W. Storti, M. A. Ganter, A. Nelson, ACS Applied Materials and Interfaces 2018, 10, 13373.

[9] T. G. Johnston, S. F. Yuan, J. M. Wagner, X. Yi, A. Saha, P. Smith, A. Nelson, H. S. Alper, Nature Communications 2020, 11, 563.

[10] T. C. Tang, E. Tham, X. Liu, K. Yehl, A. J. Rovner, H. Yuk, C. de la Fuente-Nunez, F. J. Isaacs, X. Zhao, T. K. Lu, Nature Chemical Biology 2021, 17, 724.

[11] X. Liu, T. Tang, E. Tham, H. Yuk, S. Lin, T. K. Lu, PNAS 2017, 114, DOI 10.1073/pnas.1618307114.

[12] J. L. Connell, E. T. Ritschdorff, M. Whiteley, J. B. Shear, PNAS 2013, 110, 18380.

[13] X. Liu, H. Yuk, S. Lin, G. A. Parada, T. C. Tang, E. Tham, C. de la Fuente-Nunez, T. K. Lu, X. Zhao, Advanced Materials 2018, 30, 1704821.

[14] C. Knierim, M. Enzeroth, P. Kaiser, C. Dams, D. Nette, A. Seubert, A. Klingl, C. L. Greenblatt, V. Jérôme, S. Agarwal, R. Freitag, A. Greiner, Macromolecular Bioscience 2015, 15, 1052.

[15] L. Goers, C. Ainsworth, C. H. Goey, C. Kontoravdi, P. S. Freemont, K. M. Polizzi, Biotechnology and Bioengineering 2017, 114, 1290.

[16] T. Trantidou, L. Dekker, K. Polizzi, O. Ces, Y. Elani, Interface Focus 2018, 8, DOI 10.1098/rsfs.2018.0024.

[17] I. Moya-Ramírez, P. Kotidis, M. Marbiah, J. Kim, C. Kontoravdi, K. Polizzi, ACS Synthetic Biology 2021, DOI 10.1021/acssynbio.1c00577.

[18] M. Müller, J. Becher, M. Schnabelrauch, M. Zenobi-Wong, Biofabrication 2015, 7, 035006.

[19] B. Pabst, B. Pitts, E. Lauchnor, P. S. Stewart, Antimicrobial Agents and Chemotherapy 2016, 60, 6294.

[20] M. Torres, C. Altamirano, A. J. Dickson, Current Opinion in Chemical Engineering 2018, 22, 184.

[21] J. Anderson, “Part: BBa _ J23118,” can be found under http://parts.igem.org/Part:BBa_J23118, 2006.

[22] M. Hill, M. Kern, S. Bose, A. Apostolidis, “Evaluation of an Optical CO 2 Probe for Long-Term Monitoring in Stirred-Tank Bioreactors,” can be found under https://www.presens.de/knowledge/publications/application-note/evaluation-of-an-optical-co2-probe-for-long-term-monitoring-in-stirred-tank-bioreactors-1715, **n.d.**

[23] M. Hill, G. Laslo, M. Kern, S. Bose, C. Krause, “Cell Culture Monitoring in Stirred-Tank Bioreactor with Optical pH Sensors,” can be found under https://www.presens.de/knowledge/publications/application-note/cell-culture-monitoring-in-stirred-tank-bioreactor-with-optical-ph-sensors-1717, **n.d.**

[24] M. Hill, G. Laslo, M. Kern, S. Bose, C. Krause, “Evaluation of an Optical O2 Probe and Sensor Spots for Long-Term Measurements in Stirred-Tank Bioreactors,” **n.d.**

[25] S. Iuchi, E. C. C. Lin, Proceedings of the National Academy of Sciences of the United States of America 1988, 85, 1888.

[26] M. Di Biase, P. De Leonardis, V. Castelletto, I. W. Hamley, B. Derby, N. Tirelli, Soft Matter 2011, 7, 4928.

[27] F. Yang, G. L. Nelson, Journal of Applied Polymer Science 2004, 91, 3844.

[28] A. Farrukh, W. Fan, S. Zhao, M. Salierno, J. I. Paez, A. del Campo, ChemBioChem 2018, 19, 1271.

