## Supplementary information for "Encapsulation of bacteria in bilayer Pluronic thin film hydrogels: a safe format for engineered living materials"

**Table S1: Hydrogel composition of the Plu-PluDA mixtures.** Mechanical properties of DA 0-100 hydrogels with different compositions obtained from rheology; shear storage modulus (G') of the crosslinked hydrogels.[5]

| Hydrogel Composition | Plu : PluDA | Shear storage modulus [kPa] |
|---|---|---|
| DA 0 | 100 : 0 | 17.8 ± 1.4 |
| DA 25 | 75 : 25 | 20.4 ± 2.4 |
| DA 50 | 50 : 50 | 25.9 ± 1.5 |
| DA 75 | 25 : 75 | 32.7± 3.8 |
| DA 100 | 0 : 100 | 42.9 ± 1.9 |

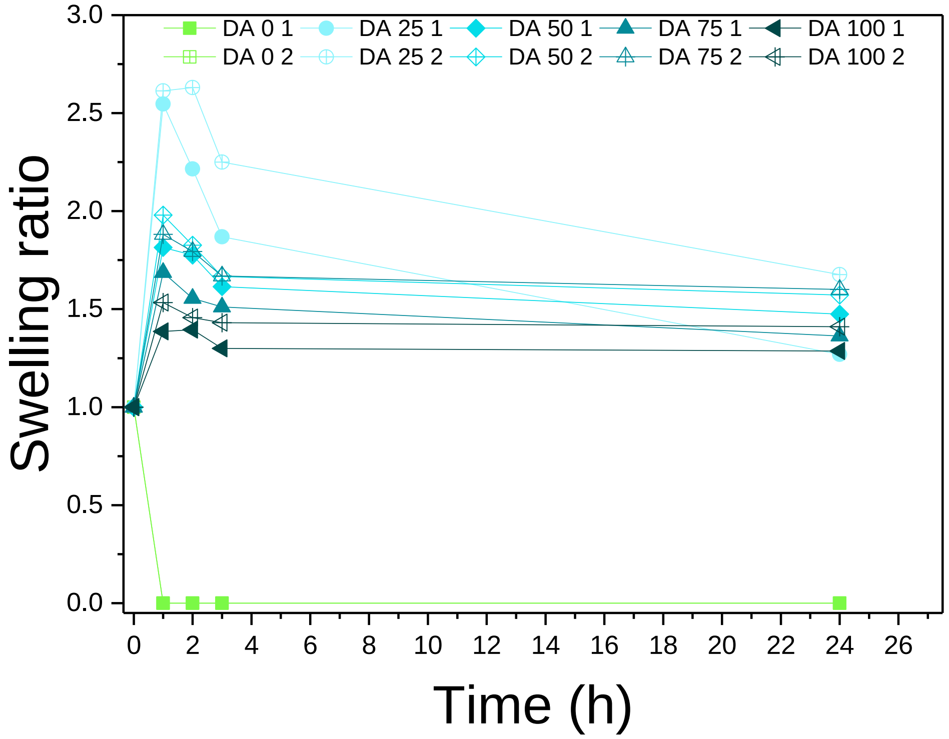


**Figure S1:** Swelling ratio (weight at time 't' compared to the initial weight) of the different DA X hydrogels in MQ water and stored at 37 °C and 5 % $CO_2$, where in DA X≥25, the ratio decreases with increasing X (i.e. covalent crosslinking) in the first hour. DA 0 dissolves in water. (n = 2)

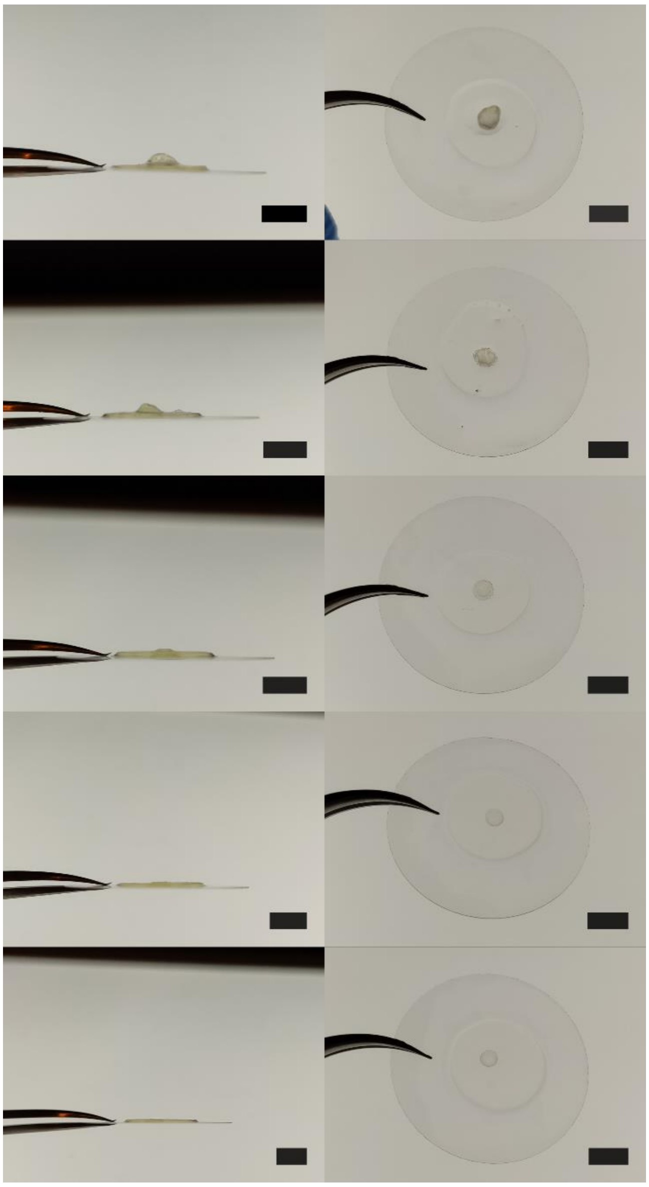

**Figure S2:** Images showing the swelling of the inner gels (DA X) of the bilayer thin films after 24 h incubation in medium at 37 °C and 5 % $CO_2$. (Scale = 5 mm)

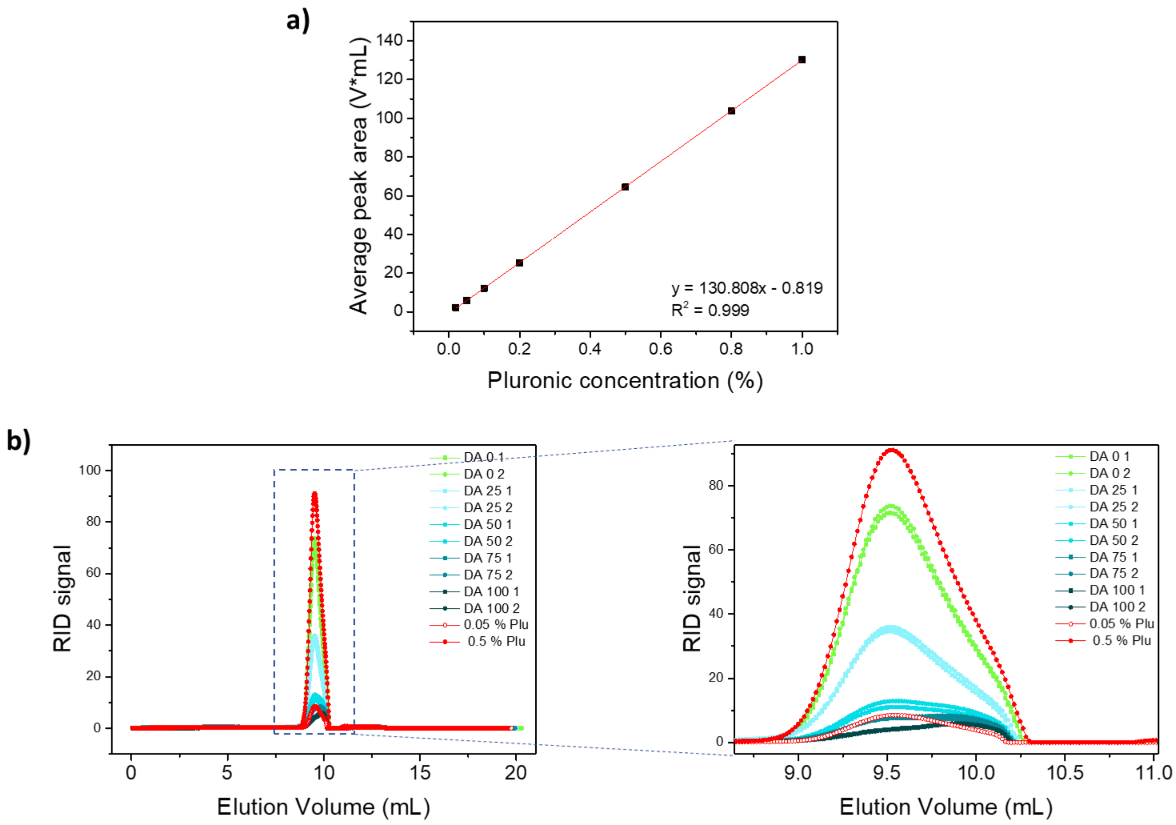


**Figure S3: a)** Standard curve with known pluronic concentrations and **b)** Representative GPC chromatograms of supernatants from DA X films after 1 h incubation. The chromatograms (in red) of solutions of known pluronic concentrations (0.05 and 0.5 % w/v) are also represented to compare the molecular weight distributions.

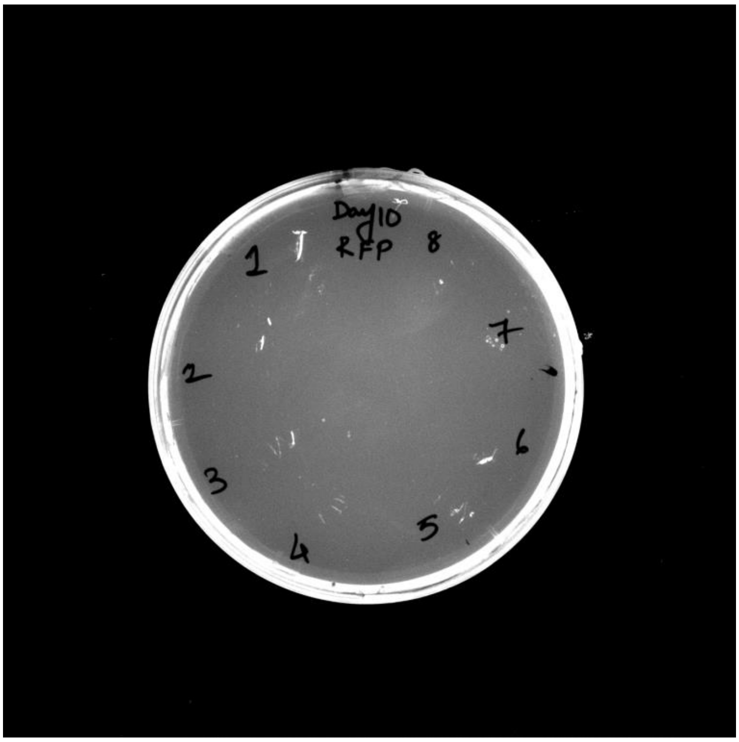


**Figure S4:** Supernatants of the thin films (N=8) embedding RFP producing bacteria for 10 days from (Figure 3d) streaked onto agar plates (made up of LB supplemented with NaCl and 50 µg/mL of Kanamycin) and incubated at 28 °C for 2 days showed no growth to confirm the absence of leakage of the thin films.

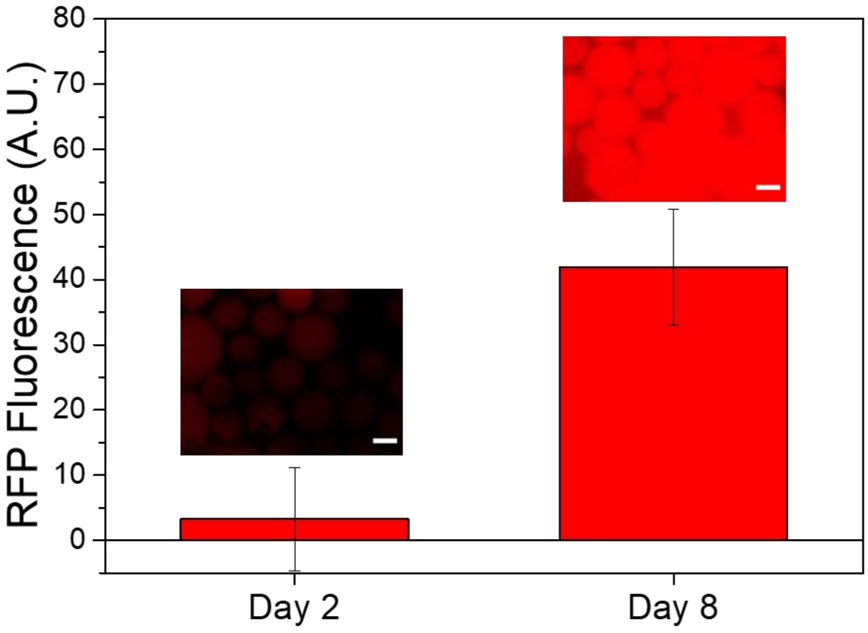


**Figure S5:** Fluorescence intensities of the Strep-Tactin affinity beads exposed to the supernatants of the thin films taken at day 2 and day 8. (scale = 50 µm)

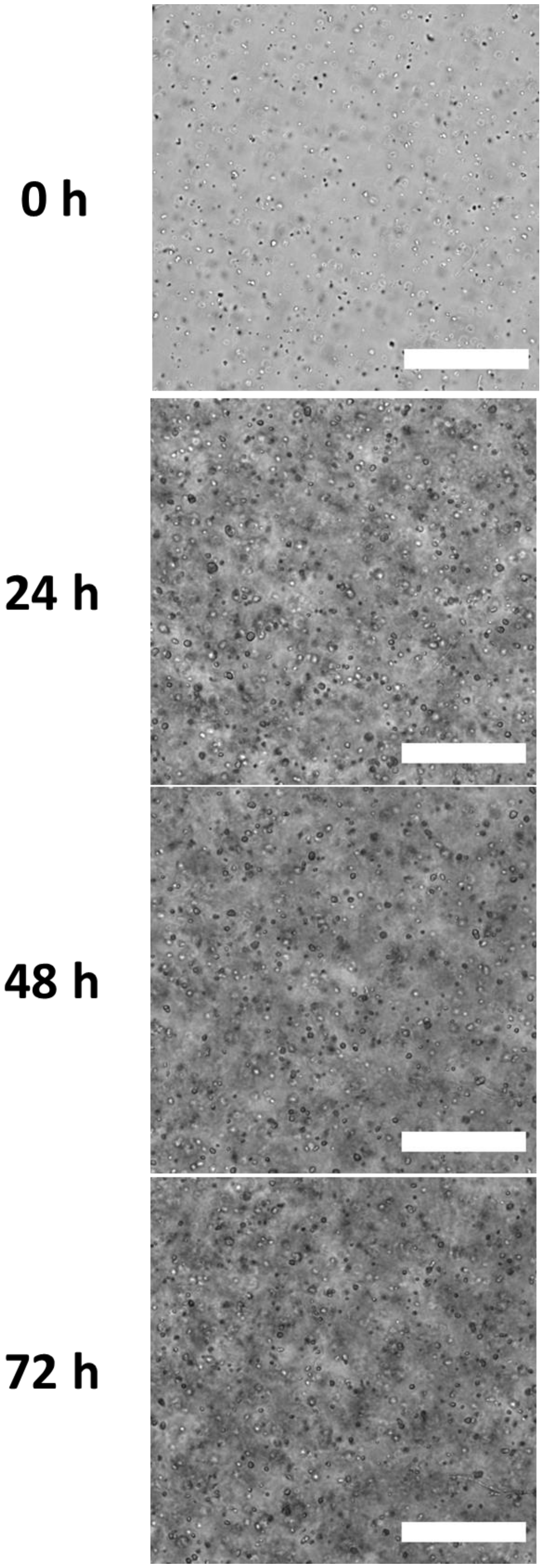


**Figure S6**: Representative images of E. coli (ClearColi) growing within the DA 50 thin film bilayers at 0, 24, 48 and 72 h. Images were taken near the middle of the bacterial gel height. (scale = 100 µm)

**a)**

**b)**

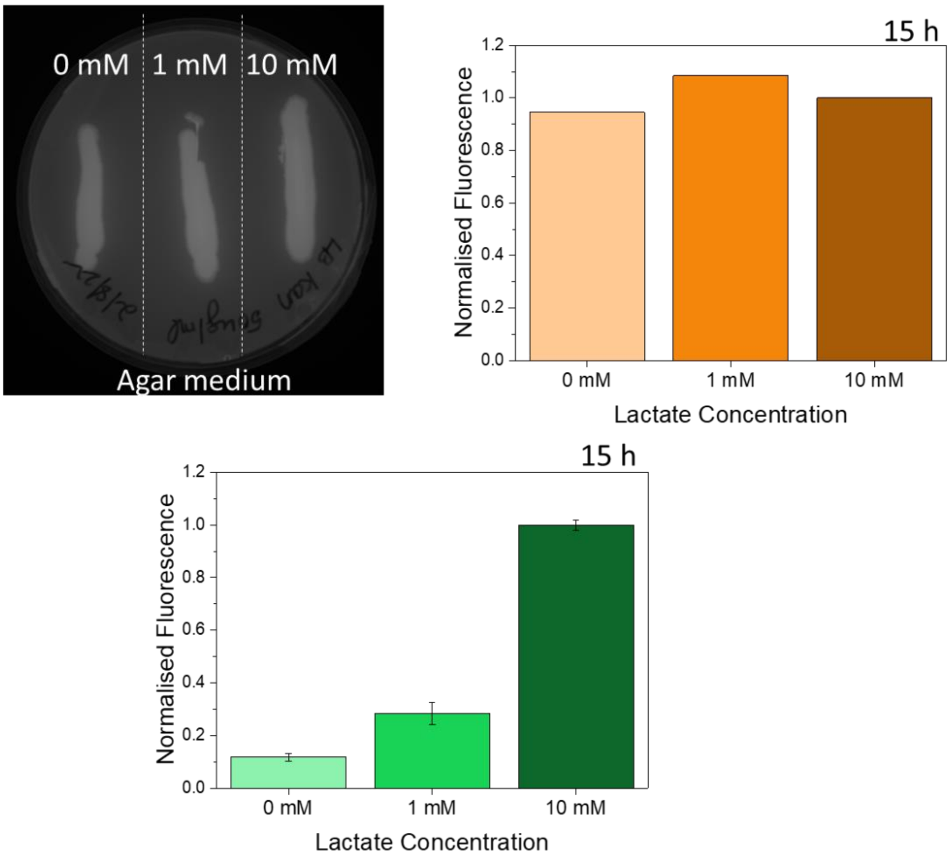


**Figure S7: a)** Fluorescence of the lactate sensing creiLOV production by the bacterial streaks on the agar plate at 15 h highlighting the strong leaky expression even in the absence of lactate (0 mM) (n = 1) and **b)** Fluorescence intensities upon lactate induction by the bacteria grown in liquid culture at 15 h. The fluorescence intensities were normalized to the corresponding 10 mM fluorescence values. (n = 2, mean ± SD)

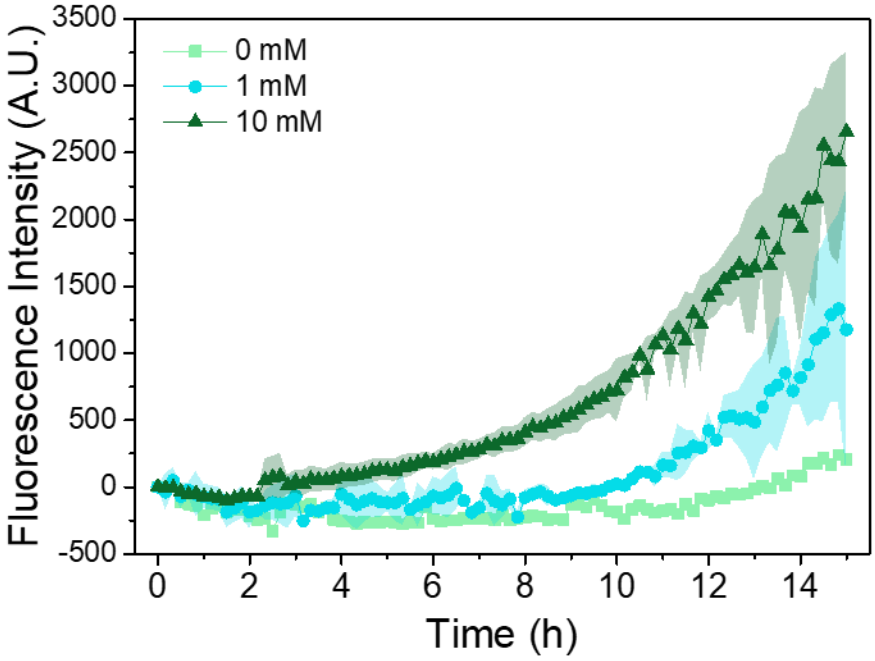


**Figure S8:** Quantification of fluorescence intensity upon lactate induction post growth for 1 day and storage in PBS for 2 days and freeze drying (N=2 for 1 and 10 mM, N=1 for 0 mM) indicating creiLOV production in the hydrogel DA 50 during 15:30 h (mean ± standard deviation).
